# Expression quantitative trait methylation analysis reveals methylomic associations with gene expression in childhood asthma

**DOI:** 10.1101/2020.02.13.937391

**Authors:** Soyeon Kim, Erick Forno, Rong Zhang, Qi Yan, Nadia Boutaoui, Edna Acosta-Pérez, Glorisa Canino, Wei Chen, Juan C. Celedón

## Abstract

Nasal airway epithelial methylation profiles have been associated with asthma, but the effects of such profiles on expression of distant cis-genes are largely unknown. We identified 16,867 significant methylation-gene expression pairs in nasal epithelium from Puerto Rican children and adolescents (with and without asthma) in an expression quantitative trait methylation (eQTM) analysis of cis-genes located within 1 Mb of the methylation probes tested. Most eQTM methylation probes were distant from their target genes, and more likely located in enhancer regions of their target genes in lung tissue than control probes. The top 500 eQTM genes were enriched in pathways for immune processes and epithelial integrity, and also more likely to be differentially expressed in atopic asthma. Moreover, we identified 5,934 paths through which methylation probes could affect atopic asthma through gene expression. Our findings suggest that distant epigenetic regulation of gene expression in airway epithelium plays a role in atopic asthma.

## Introduction

Asthma is affected not only by genetic variants but also by environmental factors such as second-hand smoke. Since DNA methylation is determined by both genetics and environment, studying methylation in relevant tissues may be key to understanding asthma pathogenesis.

A growing body of evidence suggests that abnormalities in airway epithelial integrity and function leads to interactions between injurious agents (such as pollutants and viruses) and dendritic cells, altered immune responses, and -ultimately-asthma. DNA methylation and gene expression in nasal (airway) epithelium are well correlated with those in bronchial (airway) epithelium^1^.

Because bronchial epithelial sampling requires a bronchoscopy (an invasive and costly procedure with non-trivial risks), nasal epithelial sampling is an attractive and safe approach for studies of the airway epithelium and childhood asthma^1^. Indeed, a few epigenome-wide association studies (EWAS) have identified links between DNA methylation in nasal airway epithelium and asthma. For example, we reported 7,104 CpGs associated with atopic (allergic) asthma in Puerto Ricans, an ethnic group disproportionately affected with this disease^2^.

In our prior EWAS, we estimated the effect of CpGs associated with atopic asthma on the expression of nearby genes (i.e. those adjacent to or containing a CpG of interest)^2^. More recently, we showed that most single nucleotide polymorphisms (SNPs) associated with asthma in a large meta-analysis of genome-wide association studies (GWAS) are not associated with expression of nearby genes, but rather that of more distant cis-genes within 1 Mb. Given such findings, we were interested in examining whether methylation of specific CpG sites is associated with expression of non-nearby cis-genes. We thus conducted an expression quantitative trait methylation (eQTM) analysis in nasal airway epithelium from 455 Puerto Ricans ages 9 to 20 years, including 219 subjects with asthma (cases) and 236 control subjects.

## Results

### Location of eQTM-methylation probes relative to paired genes

By testing associations between methylation probes within 1 Mb of transcription start sites (TSS) of genes and gene expression, we identified 16,867 significant methylation-expression pairs (FDR-P < 0.01, see Methods), comprising 9,103 methylation probes associated with expression of 3,512 genes. We then investigated the position of significant methylation probes in this eQTM analysis in relation to their paired genes. If a methylation probe was associated with expression of multiple genes, we counted such probe for each gene. We found that 11% and 89% of significant eQTM probes were located within and outside genes, respectively –including 4% of eQTM probes in promoter regions (**Figure 1a**). Most eQTM methylation probes were distant from their target genes (371,840 bp on average) (**Figure 1b**).

**Figure 1.**
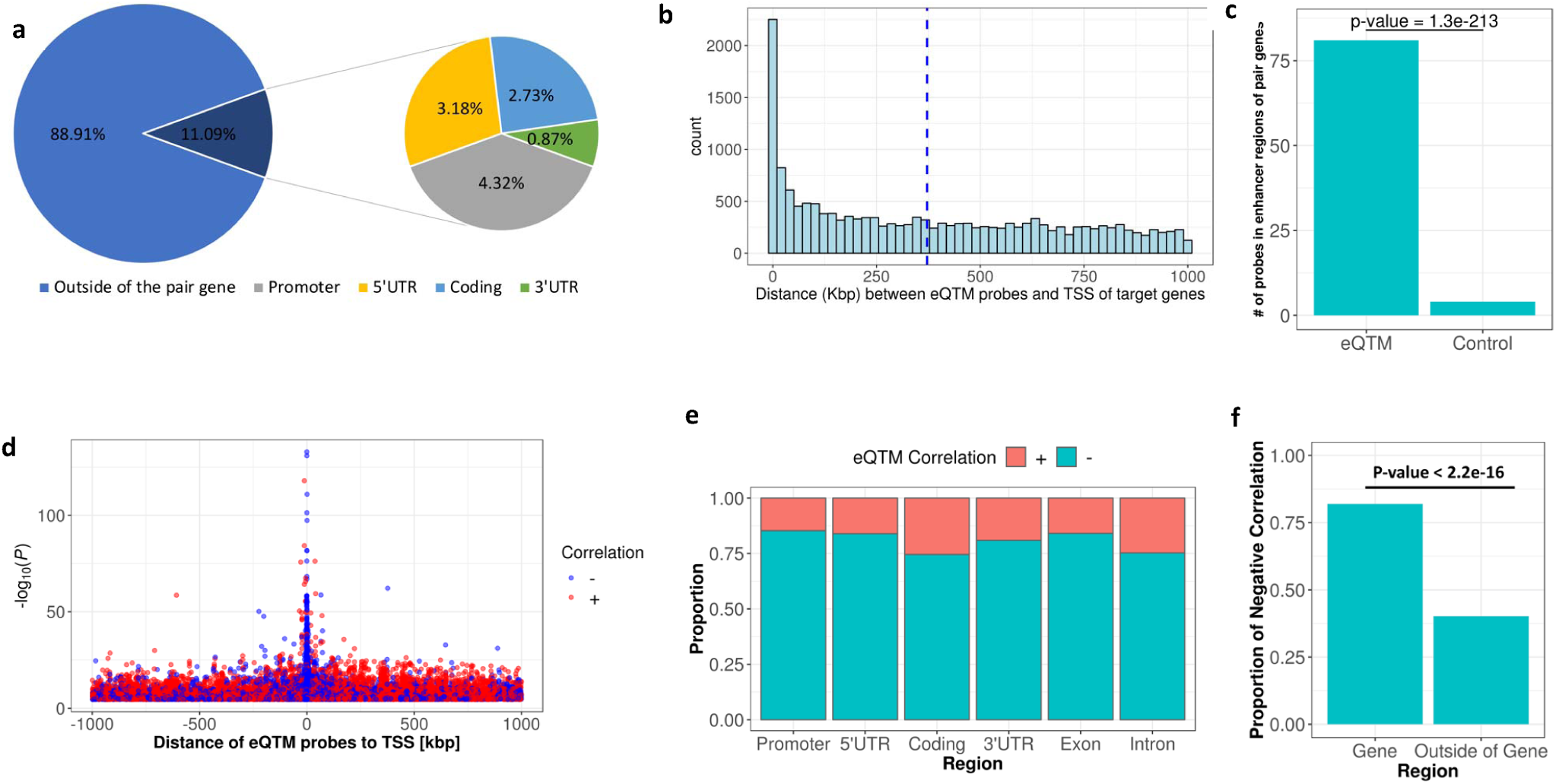
Characterization and distribution of genomic location of eQTM signals for 16,867 eQTM pairs in nasal epithelium (FDR-P < 0.01) **a.** The left chart depicts whether the probes are located inside of their paired genes. The right chart shows the specific location of probes located inside of their paired genes. **b** Distance between eQTM methylation probes and transcription start sites (TSS) of their target genes in kb pairs. **c.** Number of probes located in enhancer regions of their target genes in lung tissue. eQTM probes vs. controls (the same number of eQTM probes). Fisher’s exact test was conducted to calculate the P value. **D.** Positive/Negative correlation regarding the distance between methylation and TSS and the p-value in the eQTM analysis. **e.** The bar graph shows, within each gene region, the proportion of positive or negative correlation of the eQTM pairs. The correlation is Pearson’s correlation. **f.** The proportion of negatively correlated eQTM pairs inside (from promoter to 3’UTRs) and outside genes. The number of eQTM probes inside a gene is 1,871 and the number of the eQTM probes outside of a gene is 14,996. A chi-square test was conducted to examine the association between the region (whether the probe is located in the gene or outside of the gene) and the sign of the correlation.

Because we found distant relationships between methylation and their target genes, we assessed whether eQTM methylation probes in nasal epithelium are enriched in the enhancer regions of their paired genes. For this, we checked the enhancer database for lung tissue (http://enhanceratlas.org/)^3^, since nasal epithelial tissue was not available in that database, and nasal and bronchial epithelial methylation and expression are well correlated^1^. We found that eQTM methylation probes are more likely to be located in enhancer regions of their paired genes than randomly selected control probes (p-value: 1.3×10^−213^) (**Figure 1c, Table 1**).

**Table 1.** Top 30 eQTM methylation CpGs that are located in enhancer region of their target genes in lung tissue.

| Chr | Probe ID | Position | Gene ID | TSS<sup>a</sup> | eQTM p-value | Distance From TSS |
| --- | --- | --- | --- | --- | --- | --- |
| 16 | cg26259865 | 2880359 | *ZG16B* | 2880172 | $6.6 \times 10^{-33}$ | 187 |
| 3 | cg22012981 | 58522689 | *ACOX2* | 58522929 | $3.2 \times 10^{-31}$ | -240 |
| 3 | cg16209444 | 58522771 | *ACOX2* | 58522929 | $4.5 \times 10^{-25}$ | -158 |
| 11 | cg15453278 | 67134607 | *TBC1D10C* | 67171383 | $1.1 \times 10^{-22}$ | -36776 |
| 11 | cg21862992 | 68658383 | *MRPL21* | 68671303 | $8.6 \times 10^{-21}$ | -12920 |
| 11 | cg15453278 | 67134607 | *PTPRCAP* | 67205153 | $8.2 \times 10^{-17}$ | -70546 |
| 6 | cg25045942 | 33048291 | *HLA-DPA1* | 33041454 | $3.9 \times 10^{-15}$ | 6837 |
| 6 | cg19053046 | 33048254 | *HLA-DPA1* | 33041454 | $6.8 \times 10^{-15}$ | 6800 |
| 15 | cg10474377 | 42131658 | *JMJD7* | 42120282 | $2.1 \times 10^{-14}$ | 11376 |
| 17 | cg04204452 | 1479213 | *SERPINF2* | 1646129 | $5.1 \times 10^{-14}$ | -166916 |
| 11 | cg21920570 | 63766787 | *FERMT3* | 63974151 | $1.7 \times 10^{-13}$ | -207364 |
| 11 | cg15995296 | 67210812 | *TBC1D10C* | 67171383 | $1.5 \times 10^{-12}$ | 39429 |
| 17 | cg04204452 | 1479213 | *SERPINF1* | 1665218 | $1.5 \times 10^{-12}$ | -186005 |
| 15 | cg17163752 | 34729026 | *GOLGA8B* | 34875771 | $2.8 \times 10^{-12}$ | -146745 |
| 12 | cg21163444 | 54765670 | *NCKAP1L* | 54891494 | $1.3 \times 10^{-11}$ | -125824 |
| 11 | cg10161008 | 63766546 | *FERMT3* | 63974151 | $3.0 \times 10^{-11}$ | -207605 |
| 6 | cg17071868 | 33047056 | *HLA-DOA* | 32977389 | $1.3 \times 10^{-10}$ | 69667 |
| 11 | cg21920570 | 63766787 | *CCDC88B* | 64107689 | $3.2 \times 10^{-10}$ | -340902 |
| 3 | cg16209444 | 58522771 | *KCTD6* | 58477822 | $1.0 \times 10^{-9}$ | 44949 |
| 8 | cg20567768 | 22082066 | *SLC39A14* | 22224761 | $1.3 \times 10^{-9}$ | -142695 |
| 12 | cg22824738 | 54765988 | *NCKAP1L* | 54891494 | $2.9 \times 10^{-9}$ | -125506 |
| 11 | cg15995296 | 67210812 | *PTPRCAP* | 67205153 | $4.3 \times 10^{-9}$ | 5659 |
| 6 | cg17071868 | 33047056 | *HLA-DPB1* | 33043702 | $8.9 \times 10^{-9}$ | 3354 |
| 17 | cg01780984 | 79058859 | *BAIAP2* | 79008946 | $1.4 \times 10^{-8}$ | 49913 |
| 6 | cg19053046 | 33048254 | *HLA-DPB1* | 33043702 | $1.7 \times 10^{-8}$ | 4552 |
| 11 | cg10161008 | 63766546 | *CCDC88B* | 64107689 | $1.8 \times 10^{-8}$ | -341143 |
| 5 | cg23097826 | 149828748 | *RPS14* | 149829319 | $2.5 \times 10^{-8}$ | -571 |
| 19 | cg25264268 | 427263 | *SHC2* | 460996 | $2.5 \times 10^{-8}$ | -33733 |
| 12 | cg11700959 | 7066664 | *LAG3* | 6881669 | $3.0 \times 10^{-8}$ | 184995 |
| 12 | cg11700959 | 7066664 | *CD4* | 6898637 | $3.0 \times 10^{-8}$ | 168027 |

The eQTM analysis was conducted in nasal airway epithelium, and the enhancer regions and their target genes were identified in lung tissue (http://www.enhanceratlas.org)<sup>20</sup>. A total of 81 eQTM CpGs that are located in enhancer regions of their target genes were found. <sup>a</sup> TSS is the transcription start site of the gene.

While most methylation probes near TSS were negatively correlated with gene expression, more distant pairs tended to be positively correlated (**Figure 1d)**. Of the eQTM methylation probes associated with expression of the gene they were located in, 81.9% were negatively correlated with expression (85.3% if in promoter regions) (**Figure 1e**). In contrast, only 40.2 % of eQTM methylation probes associated with expression of a distant gene (i.e. methylation probes outside of the associated gene) were negatively correlated with expression levels (**Figure 1f)**.

Most of the top eQTM genes (by eQTM P-value) have been implicated in lung disease (**Figure 2**). *PAX8* is associated with bronchodilator response in children with asthma ^4^, *ECHDC3* is associated with obesity and asthma in children^5^, *LSP1* is associated with acute lung inflammation ^6^, *HLA-DQB1* is associated with asthma^7^ and total IgE^8^, *FRG1B* is highly mutated in lung adenocarcinoma ^9^, and *KANSL1* is associated with pulmonary function ^10^.

**Figure 2.**
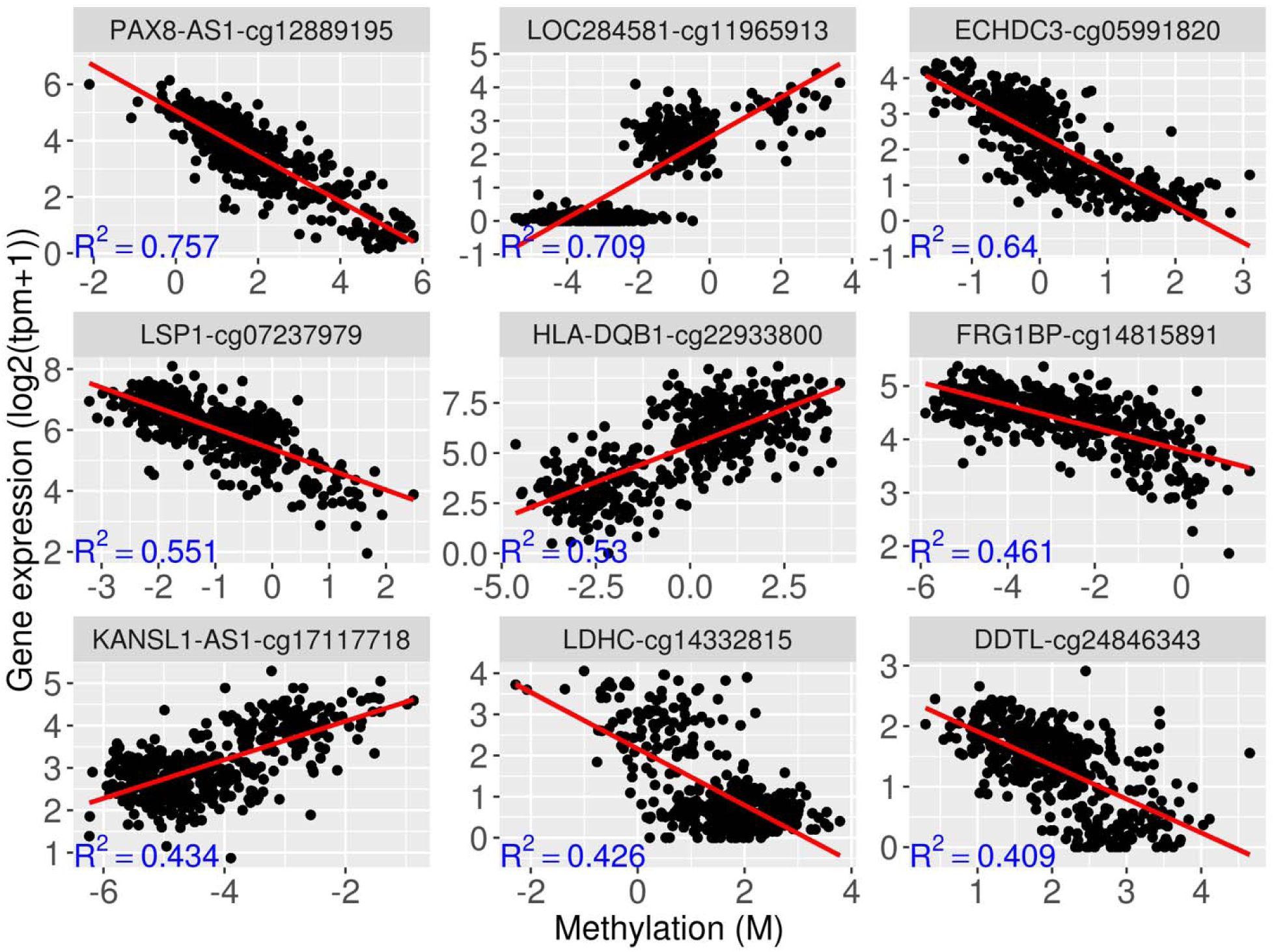
Examples of the most significantly correlated gene-methylation pairs. R^2^ is squared Pearson’s correlation between methylation and gene expression. For each gene, only the most significantly associated CpG probe is plotted.

### Gene Ontology enrichment analysis

We performed a Gene Ontology enrichment analysis including the top 500 eQTM genes. In this analysis, 34 (69.4%) of the 49 most significant gene ontology categories were related to immune processes (**Figure 3a**); the second most enriched category was cell adhesion/activation. We then investigated whether the top 500 eQTM genes are enriched for various diseases by examining significant SNPs from the GWAS catalog (https://www.ebi.ac.uk/gwas/). Most enriched diseases were related to abnormal immunity (e.g., inflammatory bowel disease and IgA nephropathy) (**Figure 3b**) and pulmonary diseases (e.g., sarcoidosis, pneumonia, and asthma).

**Figure 3.**
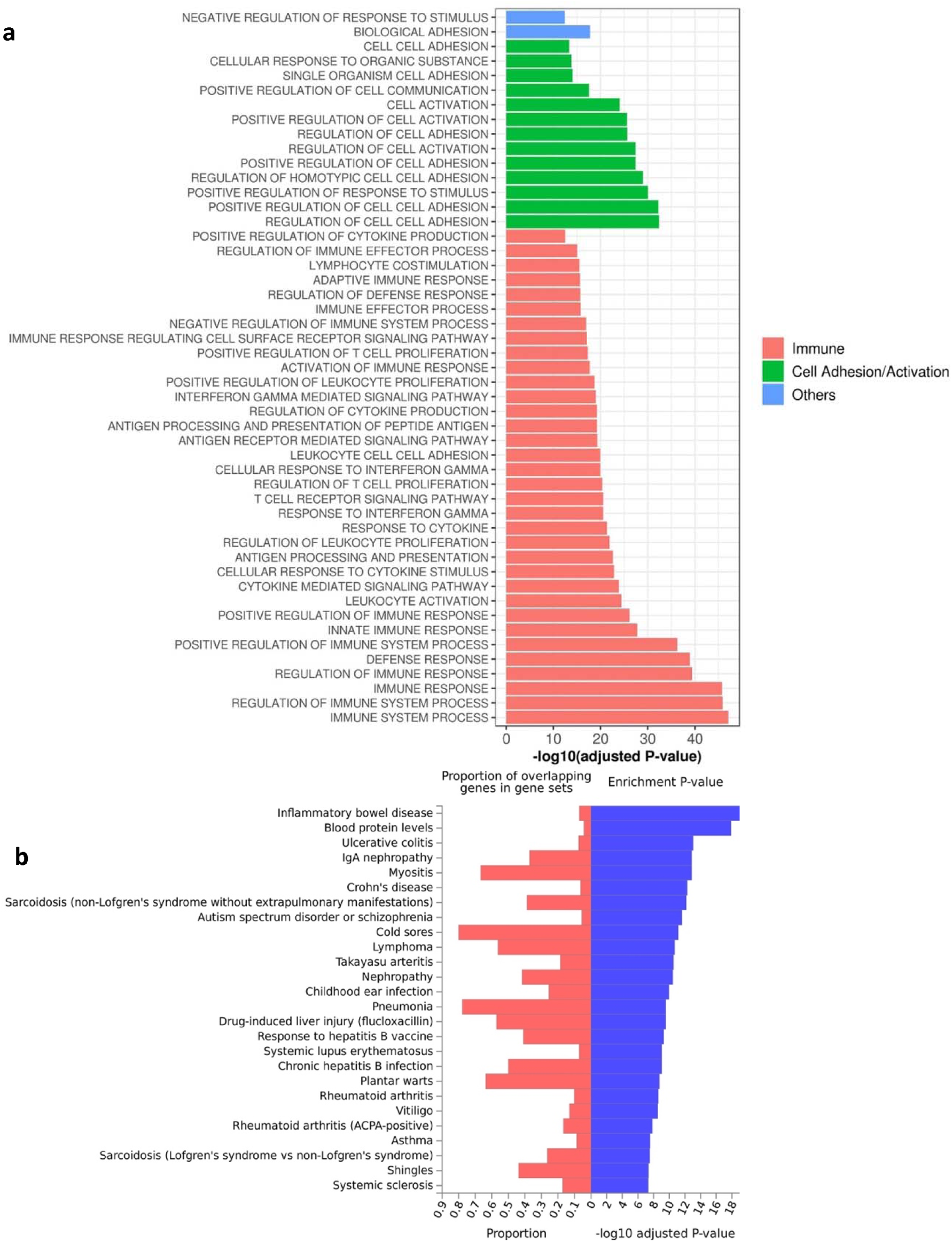
Enrichment of the top 500 eQTM genes in immune pathways/diseases. **a** Gene ontology (GO) biological processes identified for the top 500 eQTM genes. **b** Enrichment of top 500 eQTM genes among reported genes in GWAS catalog by disease. Both analyses were done through FUMA webpage^30^.

### eQTM methylation probes and genes are significantly associated with atopic asthma

We connected our eQTM results with those from our previous EWAS of atopic asthma^2^, using a genome-wide FDR-P <0.01. First, we found that only 429 (6.1%) of the 7,046 CpGs that were significantly associated with atopic asthma in our prior EWAS were associated with expression of nearby genes in the eQTM analysis (**Figure 1b**). Second, CpGs that were significant in the eQTM analysis were over-represented among CpGs that were significantly associated with atopic asthma in our prior EWAS, compared to randomly selected control CpGs (p-value < 2.2x 10^−16^) (**Figure 4a and Table 2**).

**Table 2.** Top 30 eQTM methylation probes identified in a previous epigenome-wide association study (EWAS) of atopic asthma^2^.

| probe | chr | position | EWAS<br>P-value | Associated Genes identified by eQTM (FDR < 0.01) | Nearest<br>Gene |
| --- | --- | --- | --- | --- | --- |
| cg08844313 | 5 | 149240529 | 1.2x10<sup>-16</sup> | *CD74, SLC26A2, DCTN4, AFAP1L1, GRPEL2* | *PDE6A* |
| cg20372759 | 12 | 58162287 | 1.2x10<sup>-16</sup> | *MBD6, CYP27B1, AGAP2-AS1, MARCH9, TSFM, ATP23, MYO1A* | *METTL1* |
| cg07239613 | 16 | 67051005 | 2.3x10<sup>-16</sup> | *PSMB10, FBXL8, HSF4, NOL3, CMTM4, PSKH1, TRADD, CKLF* | *CES4A* |
| cg15006973 | 1 | 35258933 | 3.2x10<sup>-16</sup> | *TMEM35B, GJB4, PSMB2, KIAA0319L, SFPQ, LOC653160* | *GJA4* |
| cg10549071 | 2 | 235160451 | 4.0x10<sup>-16</sup> | *DGKD, UGT1A1, UGT1A4, UGT1A5, UGT1A3, SCARNA5* | *SPP2* |
| cg00406211 | 10 | 121077022 | 5.6x10<sup>-16</sup> | ***GRK5***, *FAM204A, PRDX3, MCMBP* | *GRK5* |
| cg00664723 | 5 | 15927184 | 5.7x10<sup>-16</sup> | ***FBXL7*** | *FBXL7* |
| cg03875819 | 10 | 4386802 | 6.6x10<sup>-16</sup> | *AKR1C3* | *LINC00704* |
| cg21158502 | 5 | 74348187 | 8.6x10<sup>-16</sup> | *GCNT4, LINC01336, GFM2, ENC1, NSA2, FAM169A, POC5* | *ANKRD31* |
| cg13586696 | 22 | 29458723 | 1.1x10<sup>-15</sup> | *XBP1, KREMEN1, GAS2L1, RHBDD3* | *C22orf31* |
| cg20790648 | 3 | 151619923 | 1.6x10<sup>-15</sup> | *GPR171, MBNL1, AADAC, P2RY13, P2RY1* | *SUCNR1* |
| cg24707200 | 1 | 156833163 | 2.3x10<sup>-15</sup> | *SEMA4A, MEF2D, CCT3, ISG20L2, LAMTOR2, SLC25A44, RRNAD1, SMG5, UBQLN4, MRPL24* | *NTRK1* |
| cg01859321 | 8 | 144970195 | 2.5x10<sup>-15</sup> | *MROH6, SLC52A2, SCRIB, DGAT1, ZNF707, MROH1, FBXL6, PLEC, ADCK5* | *PLEC* |
| cg01870976 | 15 | 101887154 | 3.2x10<sup>-15</sup> | ***PCSK6***, *ALDH1A3, LRRK1, TM2D3* | *PCSK6* |
| cg00285620 | 11 | 102147694 | 4.9x10<sup>-15</sup> | *DYNC2H1, TMEM123, MMP10* | *BIRC3* |
| cg04132353 | 2 | 31440349 | 6.0x10<sup>-15</sup> | ***CAPN14***, *DPY30, LBH, XDH* | *CAPN14* |
| cg06675531 | 5 | 150019123 | 6.4x10<sup>-15</sup> | *ZNF300, SLC26A2, DCTN4, **SYNPO*** | *SYNPO* |
| cg22855021 | 14 | 81610812 | 7.5x10<sup>-15</sup> | *GTF2A1* | *TSHR* |
| cg09472600 | 1 | 183537770 | 9.3x10<sup>-15</sup> | *NPL, DHX9, TSEN15, APOBEC4* | *NCF2* |
| cg19107578 | 5 | 493262 | 1.2x10<sup>-14</sup> | ***SLC9A3***, *PP7080, CEP72, LOC100288152* | *SLC9A3* |
| cg18749617 | 15 | 102028637 | 1.2x10<sup>-14</sup> | ***PCSK6***, *ALDH1A3, LRRK1* | *PCSK6* |
| cg10830021 | 11 | 3815589 | 1.3x10<sup>-14</sup> | *RRM1, TRIM21, TSSC2* | *NUP98* |
| cg03387497 | 20 | 17680945 | 1.5x10<sup>-14</sup> | *POLR3F, SNRPB2, LINC00493, RRBP1, RBBP9* | *BANF2* |
| cg20337028 | 17 | 75181836 | 1.8x10<sup>-14</sup> | ***SEC14L1***, *SYNGR2, UBALD2, SEPT9, SPHK1, TNRC6C* | *SEC14L1* |
| cg17223698 | 15 | 39416631 | 2.0x10<sup>-14</sup> | *SRP14* | *C15orf54* |
| cg19497511 | 2 | 238609807 | 2.7x10<sup>-14</sup> | *COPS8* | *LRRFIP1* |
| cg08175352 | 3 | 101894206 | 3.2x10<sup>-14</sup> | *ZBTB11, TRMT10C, NXPE3, SENP7, RPL24, PCNP* | *ZPLD1* |
| cg02333649 | 22 | 19471093 | 5.5x10<sup>-14</sup> | *RTN4R, ARVCF, MRPL40, PRODH, LINC00896, UFD1L, TANGO2, DGCR8, ZDHHC8* | *CDC45* |
| cg08956463 | 6 | 41168911 | 5.9x10<sup>-14</sup> | *MDFI, TREM2, FOXP4, C6orf132, UNC5CL* | *TREML2* |
| cg04320956 | 16 | 69143512 | 6.3x10<sup>-14</sup> | ***HAS3***, *ZFP90, NQO1, NIP7, SLC7A6, CDH3, ESRP2* | *HAS3* |

Probes sorted by EWAS p-value. Genes shown in bold are the nearest gene.

**Figure 4.**
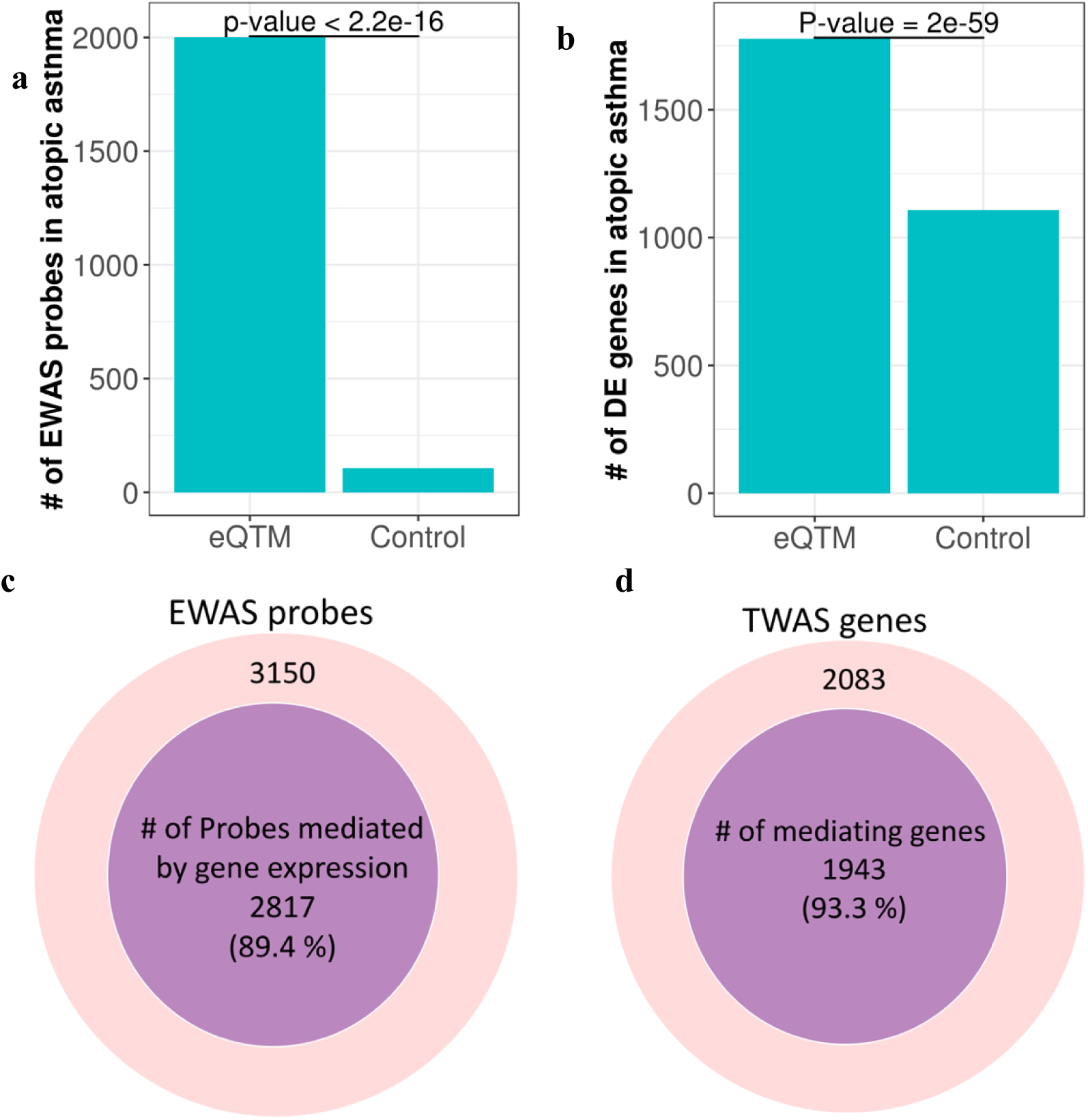
Association of eQTM methylation probes and eQTM genes with atopic asthma. **a.** Enrichment of eQTM methylation probes in epigenome-wide association studies (EWAS) of atopic asthma (genome-wide FDR-P < 0.01). eQTM refers to eQTM probes and control refers to the same number of randomly selected probes. **b.** Enrichment of eQTM genes in differentially expressed genes (DEG) in atopic asthma. DEG were identified in our previous study of EVA-PR (genome-wide FDR < 0.01)^11^. eQTM refers to eQTM genes and control refers to the same number of randomly selected genes. **c.** A majority (89.4%) of the associations between eQTM probes and atopic asthma are mediated by gene expression. **d.** A majority of (98.3%) of the genes associated with asthma mediate the association between methylation and atopic asthma.

Next, we checked whether the 3,512 significant eQTM genes identified in the current analysis are differentially expressed (DE) in atopic asthma (at genome-wide FDR-P <0.01), by checking the results of our recently published TWAS^11^. Indeed, these 3,512 eQTM genes are significantly more likely to be differentially expressed genes (DEGs) in atopic asthma than 3,512 randomly selected genes (P-value = 1.53×10^−59^) (**Figure 4b and Table 3**).

**Table 3.** Top 30 eQTM genes identified in a previous transcriptome-wide association study (TWAS) of atopic asthma^11^.

| Gene | Chr | TWAS<br>p-value | # of<br>associated<br>probes | probe | position | eQTM<br>p-value |
| --- | --- | --- | --- | --- | --- | --- |
| *CST1* | 20 | $1.1 \times 10^{-64}$ | 1 | cg14928764 | 23064608 | $5.5 \times 10^{-10}$ |
| *CLCA1* | 1 | $1.4 \times 10^{-47}$ | 5 | cg22175412 | 86063985 | $9.7 \times 10^{-15}$ |
| *NTRK2* | 9 | $4.3 \times 10^{-44}$ | 2 | cg09926027 | 87285693 | $1.5 \times 10^{-18}$ |
| *FETUB* | 3 | $7.5 \times 10^{-42}$ | 5 | cg25735294 | 186353721 | $2.6 \times 10^{-18}$ |
| *CPA3* | 3 | $3.4 \times 10^{-39}$ | 5 | cg13235059 | 149192304 | $3.1 \times 10^{-15}$ |
| *ITLN1* | 1 | $8.7 \times 10^{-38}$ | 10 | cg10094191 | 160855148 | $6.3 \times 10^{-9}$ |
| *CDH26* | 20 | $8.6 \times 10^{-37}$ | 7 | cg06943251 | 57615398 | $2.1 \times 10^{-18}$ |
| *CCL26* | 7 | $2.2 \times 10^{-35}$ | 5 | cg13053914 | 75511260 | $1.4 \times 10^{-8}$ |
| *CST2* | 20 | $4.0 \times 10^{-35}$ | 1 | cg14928764 | 23064608 | $3.2 \times 10^{-10}$ |
| *C3orf70* | 3 | $8.3 \times 10^{-35}$ | 9 | cg01390445 | 185271312 | $1.6 \times 10^{-13}$ |
| *TPSAB1* | 16 | $3.3 \times 10^{-34}$ | 26 | cg00943124 | 1705667 | $3.9 \times 10^{-14}$ |
| *CISH* | 3 | $3.0 \times 10^{-32}$ | 9 | cg23005227 | 50645426 | $8.4 \times 10^{-24}$ |
| *TPSB2* | 16 | $8.0 \times 10^{-30}$ | 20 | cg00943124 | 1705667 | $4.3 \times 10^{-13}$ |
| *ALOX15* | 17 | $3.4 \times 10^{-29}$ | 11 | cg23387401 | 4582204 | $9.7 \times 10^{-24}$ |
| *CEP72* | 5 | $1.3 \times 10^{-28}$ | 42 | cg04221910 | 616842 | $5.1 \times 10^{-21}$ |
| *SLC5A5* | 19 | $5.5 \times 10^{-28}$ | 10 | cg15734198 | 17423023 | $1.7 \times 10^{-12}$ |
| *POSTN* | 13 | $7.7 \times 10^{-28}$ | 4 | cg03071245 | 37463034 | $1.7 \times 10^{-9}$ |
| *HS3ST4* | 16 | $5.2 \times 10^{-27}$ | 4 | cg26725397 | 25937266 | $6.2 \times 10^{-15}$ |
| *PCSK6* | 15 | $1.4 \times 10^{-26}$ | 11 | cg18749617 | 102028637 | $1.4 \times 10^{-27}$ |
| *WBSCR17* | 7 | $2.5 \times 10^{-26}$ | 3 | cg01349903 | 71148142 | $5.4 \times 10^{-12}$ |
| *KYAT1* | 9 | $2.8 \times 10^{-26}$ | 12 | cg13835688 | 130859454 | $1.6 \times 10^{-16}$ |
| *ANO1* | 11 | $5.1 \times 10^{-26}$ | 7 | cg11058904 | 69987299 | $1.3 \times 10^{-15}$ |
| *ABO* | 9 | $3.0 \times 10^{-25}$ | 6 | cg11879188 | 136149908 | $7.4 \times 10^{-18}$ |
| *CMYA5* | 5 | $4.5 \times 10^{-25}$ | 1 | cg14978242 | 79501131 | $1.2 \times 10^{-7}$ |
| *SLC24A3* | 20 | $1.3 \times 10^{-23}$ | 3 | cg08371391 | 19739935 | $3.0 \times 10^{-8}$ |
| *GCNT4* | 5 | $1.1 \times 10^{-22}$ | 2 | cg21158502 | 74348187 | $2.0 \times 10^{-30}$ |
| *SLC7A1* | 13 | $1.6 \times 10^{-22}$ | 3 | cg17798847 | 30098432 | $8.3 \times 10^{-7}$ |
| *SLC45A4* | 8 | $1.8 \times 10^{-22}$ | 7 | cg07140289 | 142299684 | $1.8 \times 10^{-11}$ |
| *DQX1* | 2 | $6.9 \times 10^{-22}$ | 9 | cg02034222 | 74753281 | $4.4 \times 10^{-17}$ |
| *GSN* | 9 | $9.4 \times 10^{-22}$ | 6 | cg13928417 | 124498782 | $2.7 \times 10^{-9}$ |
| *KCNJ16* | 17 | $2.3 \times 10^{-21}$ | 2 | cg13606025 | 68070495 | $7.8 \times 10^{-10}$ |
| *LINC01336* | 5 | $7.7 \times 10^{-21}$ | 2 | cg21158502 | 74348187 | $1.6 \times 10^{-16}$ |
| *ZNF467* | 7 | $2.2 \times 10^{-20}$ | 8 | cg07970948 | 149543165 | $3.3 \times 10^{-19}$ |
| *RUSC1* | 1 | $3.7 \times 10^{-20}$ | 15 | cg23154272 | 154966068 | $2.3 \times 10^{-16}$ |
| *DHX35* | 20 | $4.7 \times 10^{-20}$ | 1 | cg26604799 | 36789861 | $2.5 \times 10^{-5}$ |
| *DPP4* | 2 | $8.1 \times 10^{-20}$ | 3 | cg22143064 | 162948592 | $5.9 \times 10^{-27}$ |
| *SOX13* | 1 | $8.4 \times 10^{-20}$ | 3 | cg17000774 | 203154457 | $1.3 \times 10^{-8}$ |
| *SLC18A2* | 10 | $1.1 \times 10^{-19}$ | 2 | cg03519180 | 119102524 | $1.2 \times 10^{-6}$ |
| *ST6GAL1* | 3 | $1.6 \times 10^{-19}$ | 6 | cg25735294 | 186353721 | $1.1 \times 10^{-20}$ |
| *C20orf197* | 20 | $3.8 \times 10^{-19}$ | 2 | cg16518142 | 58533713 | $3.9 \times 10^{-8}$ |
| *CA2* | 8 | $1.1 \times 10^{-18}$ | 1 | cg05071334 | 86195487 | $1.1 \times 10^{-5}$ |
| *NPDC1* | 9 | $2.3 \times 10^{-18}$ | 8 | cg13850871 | 139583773 | $2.6 \times 10^{-11}$ |
| *RTN4R* | 22 | $3.2 \times 10^{-18}$ | 3 | cg02333649 | 19471093 | $2.0 \times 10^{-22}$ |
| *FGF11* | 17 | $3.2 \times 10^{-18}$ | 12 | cg22637538 | 7348327 | $3.1 \times 10^{-11}$ |
| *LOC100288152* | 5 | $5.0 \times 10^{-18}$ | 15 | cg22572362 | 501938 | $1.6 \times 10^{-8}$ |
| *ELOVL5* | 6 | $5.8 \times 10^{-18}$ | 1 | cg26516974 | 52475065 | $3.5 \times 10^{-9}$ |
| *CMIP* | 16 | $1.0 \times 10^{-17}$ | 1 | cg16583186 | 81526361 | $6.7 \times 10^{-6}$ |
| *ADAMTS9* | 3 | $1.4 \times 10^{-17}$ | 11 | cg08765100 | 64211659 | $1.4 \times 10^{-11}$ |
| *CCK* | 3 | $1.8 \times 10^{-17}$ | 2 | cg07886398 | 42131702 | $1.0 \times 10^{-8}$ |

Significance for both differential expression and differential methylation defined as FDR- $P < 0.01$ . Genes shown sorted by TWAS p-value. Only the most significantly associated probe per gene is presented.

To test whether methylation affects atopic asthma through regulation of gene expression, we conducted a mediation analysis. In this analysis, we found 5,934 paths in which methylation of CpGs affect atopic asthma through gene expression, consisting of 2,817 methylation probes and 1,943 genes (**Table 4**). Of all the associations between eQTM methylation probes and atopic asthma, 89.4% were mediated by gene expression (**Figure 4c**). Likewise, 93.3% of the eQTM genes associated with atopic asthma mediate the association between methylation and atopic asthma (**Figure 4d**).

**Table 4.** Top 30 mediation paths from methylation to gene expression to atopic asthma.

| chr | probe | pos | gene | TSS | eQTM<br>p-value | EWAS<br>p-value | TWAS<br>p-value |
| --- | --- | --- | --- | --- | --- | --- | --- |
| 20 | cg14928764 | 23064608 | *CST1* | 23731574 | $5.5 \times 10^{-10}$ | $6.0 \times 10^{-4}$ | $3.2 \times 10^{-15}$ |
| 3 | cg01390445 | 185271312 | *C3orf70* | 184870802 | $1.5 \times 10^{-13}$ | $9.6 \times 10^{-8}$ | $7.0 \times 10^{-13}$ |
| 5 | cg14978242 | 79501131 | *CMYA5* | 78985658 | $1.2 \times 10^{-7}$ | $2.0 \times 10^{-7}$ | $1.4 \times 10^{-12}$ |
| 16 | cg00943124 | 1705667 | *TPSAB1* | 1290677 | $3.9 \times 10^{-14}$ | $9.0 \times 10^{-9}$ | $3.3 \times 10^{-12}$ |
| 16 | cg26725397 | 25937266 | *HS3ST4* | 25703346 | $6.2 \times 10^{-15}$ | $6.6 \times 10^{-8}$ | $6.2 \times 10^{-12}$ |
| 17 | cg23387401 | 4582204 | *ALOX15* | 4544960 | $9.7 \times 10^{-24}$ | $2.1 \times 10^{-13}$ | $6.8 \times 10^{-12}$ |
| 11 | cg11058904 | 69987299 | *ANO1* | 69924407 | $1.3 \times 10^{-15}$ | $3.9 \times 10^{-13}$ | $6.8 \times 10^{-12}$ |
| 7 | cg11303839 | 75405967 | *CCL26* | 75419064 | $7.3 \times 10^{-7}$ | $1.1 \times 10^{-7}$ | $8.9 \times 10^{-12}$ |
| 5 | cg21158502 | 74348187 | *GCNT4* | 74326724 | $2.0 \times 10^{-30}$ | $8.5 \times 10^{-16}$ | $1.1 \times 10^{-11}$ |
| 9 | cg04236137 | 123655887 | *GSN* | 124030379 | $3.7 \times 10^{-8}$ | $8.8 \times 10^{-4}$ | $1.4 \times 10^{-11}$ |
| 5 | cg21158502 | 74348187 | *LINC01336* | 74348468 | $1.6 \times 10^{-16}$ | $8.5 \times 10^{-16}$ | $1.5 \times 10^{-11}$ |
| 15 | cg18749617 | 102028637 | *PCSK6* | 102030187 | $1.4 \times 10^{-27}$ | $1.1 \times 10^{-14}$ | $1.6 \times 10^{-11}$ |
| 3 | cg13235059 | 149192304 | *CPA3* | 148583042 | $3.0 \times 10^{-15}$ | $1.7 \times 10^{-8}$ | $1.8 \times 10^{-11}$ |
| 20 | cg14928764 | 23064608 | *CST2* | 23807312 | $3.2 \times 10^{-10}$ | $6.0 \times 10^{-4}$ | $1.9 \times 10^{-11}$ |
| 5 | cg01181940 | 478916 | *CEP72* | 612404 | $1.6 \times 10^{-20}$ | $2.2 \times 10^{-11}$ | $2.8 \times 10^{-11}$ |
| 9 | cg09926027 | 87285693 | *NTRK2* | 87283372 | $1.5 \times 10^{-18}$ | $4.5 \times 10^{-12}$ | $3.2 \times 10^{-11}$ |
| 1 | cg23154272 | 154966068 | *RUSC1* | 155290639 | $2.2 \times 10^{-16}$ | $2.0 \times 10^{-8}$ | $3.4 \times 10^{-11}$ |
| 9 | cg11879188 | 136149908 | *ABO* | 136150630 | $7.3 \times 10^{-18}$ | $1.9 \times 10^{-7}$ | $3.8 \times 10^{-11}$ |
| 3 | cg23005227 | 50645426 | *CISH* | 50649262 | $8.3 \times 10^{-24}$ | $4.0 \times 10^{-12}$ | $3.8 \times 10^{-11}$ |
| 6 | cg14178895 | 11778902 | *ADTRP* | 11779280 | $6.8 \times 10^{-23}$ | $3.6 \times 10^{-8}$ | $4.0 \times 10^{-11}$ |
| 3 | cg25735294 | 186353721 | *ST6GAL1* | 186648314 | $1.0 \times 10^{-20}$ | $3.3 \times 10^{-10}$ | $5.2 \times 10^{-11}$ |
| 8 | cg07140289 | 142299684 | *SLC45A4* | 142238673 | $1.8 \times 10^{-11}$ | $2.4 \times 10^{-4}$ | $6.3 \times 10^{-11}$ |
| 2 | cg04132353 | 31440349 | *CAPN14* | 31440411 | $4.6 \times 10^{-20}$ | $5.9 \times 10^{-15}$ | $6.4 \times 10^{-11}$ |
| 5 | cg14978242 | 79501131 | *SERINC5* | 79551901 | $1.3 \times 10^{-9}$ | $2.0 \times 10^{-7}$ | $7.7 \times 10^{-11}$ |
| 1 | cg03058346 | 91275170 | *LRRC8D* | 90286572 | $1.4 \times 10^{-6}$ | $6.9 \times 10^{-4}$ | $8.4 \times 10^{-11}$ |
| 1 | cg01062020 | 162382848 | *SH2D1B* | 162381928 | $1.7 \times 10^{-6}$ | $2.0 \times 10^{-6}$ | $9.8 \times 10^{-11}$ |
| 20 | cg26604799 | 36789861 | *DHX35* | 37590980 | $2.5 \times 10^{-5}$ | $2.5 \times 10^{-3}$ | $1.0 \times 10^{-10}$ |
| 16 | cg00943124 | 1705667 | *TPSB2* | 1280185 | $4.2 \times 10^{-13}$ | $9.0 \times 10^{-9}$ | $1.1 \times 10^{-10}$ |
| 2 | cg22143064 | 162948592 | *DPP4* | 162931052 | $5.8 \times 10^{-27}$ | $5.2 \times 10^{-8}$ | $1.5 \times 10^{-10}$ |
| 15 | cg09407660 | 59910436 | *GCNT3* | 59903981 | $1.3 \times 10^{-19}$ | $9.8 \times 10^{-10}$ | $1.5 \times 10^{-10}$ |

Results sorted by TWAS p-value<sup>11</sup>. Total of 5394 mediation paths were identified. Only one mediation path was presented per gene. Mediation analysis was conducted using Baron and Kenny (1986).

### eQTM results in EVA-PR are replicated in an African American cohort

To attempt replication of our eQTM results in EVA-PR, we used public data from GSE65205^12^, which includes both methylation and gene expression array data in nasal epithelium for 69 children (36 with atopic asthma and 33 healthy controls, mostly [91.3%] African American). Using a similar approach to that used in EVA-PR, this replication eQTM analysis was adjusted for age, sex, race/ethnicity, atopic asthma status, and unobserved batch effects.

Of the 16,867 significant associations between methylation and gene expression in EVA-PR, we were able to test 14,397 associations in GSE65205, due to differences in the platforms used to assess gene expression (RNA-Seq vs. microarray). Of these 14,397 methylation-expression pairs, 12,559 (87.2 %) had the same direction of association in GSE65205. Despite the small sample size of GSE65205, 6,562 (45.6%) of the significant associations in EVA-PR were replicated at FDR-P < 0.05, in the same direction of association (**Table 5**). These replicated associations include 3,992 methylation probes and 1,106 genes. Of the 3,992 replicated methylation probes, 3,222 probes were tested in our prior EWAS in EVA-PR^2^: 1,412 (43.8%) of these 3,222 probes are significantly associated with atopic asthma (FDR-P < 0.05) (**Table 6**).

**Table 5.** Top 30 most significant eQTM methylation-gene pairs in EVA-PR cohort that replicated in GSE65205.

| Associated pairs |  | EVA-PR |  |  | GSE65205 |  |  |
| --- | --- | --- | --- | --- | --- | --- | --- |
| probe | gene | beta | p-value | FDR | beta | p-value | FDR |
| cg05991820 | *ECHDC3* | -1 | $5.1 \times 10^{-98}$ | $7.2 \times 10^{-92}$ | -0.31 | $3.9 \times 10^{-3}$ | $1.2 \times 10^{-2}$ |
| cg22933800 | *HLA-DQB1* | 0.75 | $1.7 \times 10^{-76}$ | $1.2 \times 10^{-70}$ | 1 | $7.9 \times 10^{-20}$ | $5.7 \times 10^{-16}$ |
| cg07237979 | *LSP1* | -0.67 | $4.6 \times 10^{-69}$ | $3.0 \times 10^{-63}$ | -0.57 | $2.4 \times 10^{-12}$ | $4.9 \times 10^{-10}$ |
| cg14332815 | *LDHC* | -0.71 | $1.3 \times 10^{-56}$ | $4.0 \times 10^{-51}$ | -1.1 | $1.5 \times 10^{-18}$ | $5.3 \times 10^{-15}$ |
| cg10296238 | *SPATC1L* | -0.31 | $1.1 \times 10^{-51}$ | $2.5 \times 10^{-46}$ | -0.23 | $2.4 \times 10^{-10}$ | $1.7 \times 10^{-8}$ |
| cg17117718 | *CRHR1-IT1* | 0.29 | $4.0 \times 10^{-51}$ | $8.6 \times 10^{-46}$ | 0.35 | $5.2 \times 10^{-8}$ | $1.2 \times 10^{-6}$ |
| cg16145915 | *ZFAND2A* | 0.48 | $2.6 \times 10^{-50}$ | $5.1 \times 10^{-45}$ | 0.53 | $1.1 \times 10^{-4}$ | $6.0 \times 10^{-4}$ |
| cg10626236 | *CDK11A* | 0.59 | $3.2 \times 10^{-50}$ | $6.2 \times 10^{-45}$ | 0.39 | $1.0 \times 10^{-2}$ | $2.5 \times 10^{-2}$ |
| cg03190825 | *CYP4F11* | -1.3 | $2.5 \times 10^{-48}$ | $4.5 \times 10^{-43}$ | -0.41 | $6.7 \times 10^{-3}$ | $1.8 \times 10^{-2}$ |
| cg22092521 | *CFD* | -0.83 | $1.7 \times 10^{-45}$ | $2.6 \times 10^{-40}$ | -1.2 | $1.2 \times 10^{-6}$ | $1.4 \times 10^{-5}$ |
| cg11375102 | *TMEM204* | -0.68 | $1.3 \times 10^{-44}$ | $1.9 \times 10^{-39}$ | -1 | $1.7 \times 10^{-14}$ | $1.2 \times 10^{-11}$ |
| cg24977027 | *THNSL2* | -0.78 | $5.6 \times 10^{-44}$ | $8.0 \times 10^{-39}$ | -1.2 | $4.6 \times 10^{-13}$ | $1.3 \times 10^{-10}$ |
| cg05461841 | *ZG16B* | -0.66 | $2.0 \times 10^{-42}$ | $2.6 \times 10^{-37}$ | -0.71 | $1.4 \times 10^{-4}$ | $7.2 \times 10^{-4}$ |
| cg06851207 | *PNMAL1* | -0.79 | $2.1 \times 10^{-42}$ | $2.7 \times 10^{-37}$ | -0.89 | $3.0 \times 10^{-7}$ | $4.6 \times 10^{-6}$ |
| cg01878807 | *DHRS4-AS1* | -0.42 | $4.3 \times 10^{-42}$ | $5.4 \times 10^{-37}$ | -0.34 | $1.3 \times 10^{-5}$ | $1.0 \times 10^{-4}$ |
| cg02926397 | *LY6D* | -1.2 | $8.3 \times 10^{-42}$ | $1.0 \times 10^{-36}$ | -2.5 | $6.6 \times 10^{-10}$ | $3.8 \times 10^{-8}$ |
| cg02719634 | *SLC22A18AS* | -0.28 | $5.8 \times 10^{-41}$ | $6.6 \times 10^{-36}$ | -0.41 | $3.0 \times 10^{-6}$ | $3.0 \times 10^{-5}$ |
| cg06846259 | *POMC* | -0.5 | $5.9 \times 10^{-41}$ | $6.6 \times 10^{-36}$ | -1 | $1.2 \times 10^{-16}$ | $2.4 \times 10^{-13}$ |
| cg15176213 | *COX7A1* | -0.85 | $6.3 \times 10^{-41}$ | $6.9 \times 10^{-36}$ | -1.3 | $2.4 \times 10^{-14}$ | $1.7 \times 10^{-11}$ |
| cg14815891 | *FRG1BP* | -0.19 | $8.0 \times 10^{-41}$ | $8.7 \times 10^{-36}$ | -0.26 | $1.3 \times 10^{-4}$ | $7.0 \times 10^{-4}$ |
| cg24846343 | *GSTT2B* | -0.79 | $4.2 \times 10^{-39}$ | $4.2 \times 10^{-34}$ | -1 | $1.1 \times 10^{-12}$ | $2.6 \times 10^{-10}$ |
| cg19059861 | *BPIFA1* | -3.2 | $4.6 \times 10^{-37}$ | $4.1 \times 10^{-32}$ | -3.9 | $1.4 \times 10^{-8}$ | $4.1 \times 10^{-7}$ |
| cg22933800 | *HLA-DQA2* | -0.49 | $9.0 \times 10^{-37}$ | $7.8 \times 10^{-32}$ | -0.18 | $1.7 \times 10^{-5}$ | $1.3 \times 10^{-4}$ |
| cg06322601 | *RASA4* | 0.18 | $3.8 \times 10^{-35}$ | $3.0 \times 10^{-30}$ | 0.08 | $1.6 \times 10^{-2}$ | $3.7 \times 10^{-2}$ |
| cg05681977 | *SLC39A4* | -0.37 | $6.4 \times 10^{-34}$ | $4.7 \times 10^{-29}$ | -0.64 | $9.1 \times 10^{-16}$ | $1.4 \times 10^{-12}$ |
| cg10207745 | *LINC01559* | -0.86 | $7.3 \times 10^{-34}$ | $5.2 \times 10^{-29}$ | -0.52 | $3.0 \times 10^{-3}$ | $9.5 \times 10^{-3}$ |
| cg10807101 | *GSTM3* | -0.43 | $3.4 \times 10^{-33}$ | $2.3 \times 10^{-28}$ | -0.68 | $4.9 \times 10^{-7}$ | $6.8 \times 10^{-6}$ |
| cg08450017 | *CXCR6* | -0.46 | $2.0 \times 10^{-32}$ | $1.3 \times 10^{-27}$ | -0.59 | $2.4 \times 10^{-10}$ | $1.7 \times 10^{-8}$ |
| cg01850135 | *NLRC3* | -0.34 | $8.3 \times 10^{-32}$ | $5.2 \times 10^{-27}$ | -0.9 | $1.5 \times 10^{-11}$ | $2.0 \times 10^{-9}$ |
| cg23161218 | *ACAP1* | 0.31 | $1.2 \times 10^{-31}$ | $7.5 \times 10^{-27}$ | 0.29 | $2.1 \times 10^{-4}$ | $1.0 \times 10^{-3}$ |

Replication defined as FDR $P < 0.05$ with effect in the same direction as in EVA-PR.

**Table 6.** Top 30 most significant eQTM methylation-gene pairs in EVA-PR cohort that replicated in GSE65205. Only eQTM probes that are associated with atopic asthma in EVA-PR cohort (FDR-P < 0.05) are presented.

| Associated pairs |  | EVA-PR |  |  | GSE65205 |  |  | EWAS<br>(EVA-PR) |
| --- | --- | --- | --- | --- | --- | --- | --- | --- |
| probe | gene | beta | p-value | FDR | beta | p-value | FDR | FDR |
| cg04511125 | *THNSL2* | -0.9 | $1.1 \times 10^{-36}$ | $9.2 \times 10^{-32}$ | -0.81 | $1.2 \times 10^{-6}$ | $1.4 \times 10^{-5}$ | $7.3 \times 10^{-3}$ |
| cg17252645 | *LY6D* | -1 | $6.1 \times 10^{-36}$ | $5.0 \times 10^{-31}$ | -1.6 | $1.5 \times 10^{-8}$ | $4.4 \times 10^{-7}$ | $4.1 \times 10^{-2}$ |
| cg10807101 | *GSTM3* | -0.43 | $3.4 \times 10^{-33}$ | $2.3 \times 10^{-28}$ | -0.68 | $4.9 \times 10^{-7}$ | $6.8 \times 10^{-6}$ | $7.6 \times 10^{-4}$ |
| cg08450017 | *CXCR6* | -0.46 | $2.0 \times 10^{-32}$ | $1.3 \times 10^{-27}$ | -0.59 | $2.4 \times 10^{-10}$ | $1.7 \times 10^{-8}$ | $2.6 \times 10^{-4}$ |
| cg01850135 | *NLRC3* | -0.34 | $8.3 \times 10^{-32}$ | $5.2 \times 10^{-27}$ | -0.9 | $1.5 \times 10^{-11}$ | $2.0 \times 10^{-9}$ | $5.6 \times 10^{-4}$ |
| cg23161218 | *ACAP1* | 0.31 | $1.2 \times 10^{-31}$ | $7.5 \times 10^{-27}$ | 0.29 | $2.1 \times 10^{-4}$ | $1.0 \times 10^{-3}$ | $2.3 \times 10^{-3}$ |
| cg22012981 | *ACOX2* | -0.86 | $3.2 \times 10^{-31}$ | $1.9 \times 10^{-26}$ | -0.35 | $1.1 \times 10^{-2}$ | $2.6 \times 10^{-2}$ | $3.3 \times 10^{-8}$ |
| cg07786657 | *CD247* | -0.35 | $4.6 \times 10^{-31}$ | $2.7 \times 10^{-26}$ | -0.37 | $3.9 \times 10^{-9}$ | $1.5 \times 10^{-7}$ | $8.0 \times 10^{-4}$ |
| cg03546687 | *IL32* | 0.59 | $2.4 \times 10^{-29}$ | $1.3 \times 10^{-24}$ | 0.75 | $5.0 \times 10^{-6}$ | $4.5 \times 10^{-5}$ | $3.1 \times 10^{-3}$ |
| cg14527029 | *HGD* | -0.76 | $3.2 \times 10^{-29}$ | $1.6 \times 10^{-24}$ | -1.1 | $9.2 \times 10^{-8}$ | $1.8 \times 10^{-6}$ | $1.4 \times 10^{-7}$ |
| cg11453837 | *PSMB9* | -0.98 | $3.3 \times 10^{-28}$ | $1.6 \times 10^{-23}$ | -0.78 | $6.0 \times 10^{-4}$ | $2.5 \times 10^{-3}$ | $1.7 \times 10^{-4}$ |
| cg27583010 | *SEPT1* | -0.39 | $1.0 \times 10^{-27}$ | $4.6 \times 10^{-23}$ | -0.61 | $3.4 \times 10^{-9}$ | $1.3 \times 10^{-7}$ | $4.3 \times 10^{-2}$ |
| cg18749617 | *PCSK6* | -0.38 | $1.4 \times 10^{-27}$ | $6.3 \times 10^{-23}$ | -0.22 | $1.1 \times 10^{-3}$ | $4.1 \times 10^{-3}$ | $1.2 \times 10^{-10}$ |
| cg08450017 | *CCR5* | -0.36 | $2.8 \times 10^{-27}$ | $1.2 \times 10^{-22}$ | -0.54 | $3.3 \times 10^{-9}$ | $1.3 \times 10^{-7}$ | $2.6 \times 10^{-4}$ |
| cg22143064 | *DPP4* | -1.1 | $5.9 \times 10^{-27}$ | $2.4 \times 10^{-22}$ | -0.87 | $4.7 \times 10^{-9}$ | $1.8 \times 10^{-7}$ | $1.8 \times 10^{-5}$ |
| cg07786657 | *RCSD1* | -0.27 | $3.2 \times 10^{-26}$ | $1.2 \times 10^{-21}$ | -0.39 | $2.0 \times 10^{-5}$ | $1.4 \times 10^{-4}$ | $8.0 \times 10^{-4}$ |
| cg08159663 | *NLRC5* | -0.63 | $1.3 \times 10^{-25}$ | $4.6 \times 10^{-21}$ | -0.31 | $1.2 \times 10^{-2}$ | $2.9 \times 10^{-2}$ | $1.3 \times 10^{-4}$ |
| cg19517476 | *RASAL3* | -0.27 | $2.6 \times 10^{-25}$ | $9.0 \times 10^{-21}$ | -0.87 | $1.6 \times 10^{-13}$ | $6.5 \times 10^{-11}$ | $2.8 \times 10^{-4}$ |
| cg12044599 | *TBC1D10C* | -0.32 | $3.5 \times 10^{-25}$ | $1.2 \times 10^{-20}$ | -0.71 | $3.6 \times 10^{-7}$ | $5.4 \times 10^{-6}$ | $3.5 \times 10^{-3}$ |
| cg26833120 | *LCK* | 0.46 | $4.8 \times 10^{-25}$ | $1.6 \times 10^{-20}$ | 0.73 | $4.8 \times 10^{-5}$ | $3.1 \times 10^{-4}$ | $1.6 \times 10^{-3}$ |
| cg02297541 | *HLA-DMA* | 0.49 | $5.3 \times 10^{-25}$ | $1.8 \times 10^{-20}$ | 0.91 | $6.6 \times 10^{-10}$ | $3.7 \times 10^{-8}$ | $9.2 \times 10^{-4}$ |
| cg06148175 | *ACY3* | -0.65 | $5.7 \times 10^{-25}$ | $1.9 \times 10^{-20}$ | -0.35 | $2.8 \times 10^{-3}$ | $8.9 \times 10^{-3}$ | $2.7 \times 10^{-2}$ |
| cg00676801 | *STAT1* | -0.55 | $7.0 \times 10^{-25}$ | $2.2 \times 10^{-20}$ | -1.3 | $8.2 \times 10^{-8}$ | $1.7 \times 10^{-6}$ | $8.5 \times 10^{-4}$ |
| cg12911952 | *SLC22A18AS* | -0.39 | $1.2 \times 10^{-24}$ | $3.8 \times 10^{-20}$ | -1.1 | $2.7 \times 10^{-7}$ | $4.3 \times 10^{-6}$ | $9.4 \times 10^{-5}$ |
| cg05141234 | *HLA-DMB* | 0.57 | $1.8 \times 10^{-24}$ | $5.3 \times 10^{-20}$ | 0.75 | $3.5 \times 10^{-7}$ | $5.3 \times 10^{-6}$ | $2.9 \times 10^{-4}$ |
| cg00945209 | *TMC8* | -0.54 | $2.2 \times 10^{-24}$ | $6.5 \times 10^{-20}$ | -1.1 | $1.8 \times 10^{-9}$ | $8.2 \times 10^{-8}$ | $1.7 \times 10^{-2}$ |
| cg09878888 | *LPXN* | 0.24 | $3.2 \times 10^{-24}$ | $9.1 \times 10^{-20}$ | 0.35 | $6.7 \times 10^{-3}$ | $1.8 \times 10^{-2}$ | $1.5 \times 10^{-4}$ |
| cg00676801 | *STAT4* | -0.33 | $3.4 \times 10^{-24}$ | $9.7 \times 10^{-20}$ | -0.57 | $2.0 \times 10^{-8}$ | $5.5 \times 10^{-7}$ | $8.5 \times 10^{-4}$ |
| cg23387401 | *ALOX15* | -0.37 | $9.7 \times 10^{-24}$ | $2.6 \times 10^{-19}$ | -0.64 | $6.2 \times 10^{-8}$ | $1.4 \times 10^{-6}$ | $1.0 \times 10^{-9}$ |
| cg13443575 | *CCL5* | 0.54 | $1.3 \times 10^{-23}$ | $3.5 \times 10^{-19}$ | 0.77 | $7.8 \times 10^{-6}$ | $6.6 \times 10^{-5}$ | $5.5 \times 10^{-3}$ |

Replication defined as FDR P<0.05 with effect in the same direction as in EVA-PR. Only one gene per methylation probe presented.

## Discussion

To date, there have been much fewer eQTM studies than eQTL studies^13^, despite probable large joint causal effects of DNA methylation and gene expression on complex diseases. While genotype does not change as a disease progresses, both epigenetic regulation and transcriptomic activity change as a disease develops or worsens. Thus, studying eQTM may complement findings from genetic or eQTL studies and add novel insights into disease pathogenesis.

Most previous genome-wide eQTM studies have been limited to healthy subjects^14,15^. In the few instances in which both subjects with asthma and healthy controls were included, only CpGs that were significant in an EWAS –and only genes nearby those CpGs (e.g., within 10 kb)– were examined 16-18. In contrast, we assessed all genome-wide CpGs along with expression of cis-genes located within 1 Mb in the current analysis of children and adolescents with and without asthma. Moreover, we were able to replicate nearly half of our significant findings in an independent cohort of predominantly African American children.

Notably, in our analysis most significant eQTM methylation probes were not nearby their target cis-genes, a finding that may be explained by physical contact between CpG sites and promoter/coding regions of distant target genes through looping chromatin structures^19^. Significant eQTM probes were also more likely to be localized in enhancer regions of their target genes in lung tissue than control probes, suggesting that CpG sites can affect transcription of non-nearby (distant) cis-genes through enhancer activity. We also found that while most methylation probes near TSS were negatively correlated with gene expression, more distant pairs tended to be positively correlated. Consistent with our findings, methylation in promoter regions and the first intron have been negatively correlated with gene expression, while methylation of more distant CpG sites and gene expression has been positively correlated with gene expression in several types of cancer^20,21 22^.

We show an over-representation of the top eQTM methylation probes among CpGs associated with atopic asthma. Similarly, we report an over-representation of the top eQTM genes among DEG in atopic asthma. Moreover, we show that most associations between eQTM methylation probes and atopic asthma are mediated by gene expression. Given that we also found that eQTM methylation probes in nasal epithelium are over-represented in enhancer regions of their paired genes in lung tissue, our findings provide further support for studies of nasal epithelial epigenomics and transcriptomics as a valid alternative to more invasive and costly studies of bronchial epithelium.

We recognize several study limitations. First, we only included subjects in a high-risk population (Puerto Ricans). However, we have previously replicated findings from GWAS^9^ and EWAS ^11^ of asthma in Puerto Ricans in other racial or ethnic groups, including non-Hispanic whites, African Americans, and members of other Hispanic subgroups. Moreover, about half of the significant eQTM pairs in the current analysis in Puerto Ricans were significant in African Americans, despite the small sample size of the replication cohort. Second, we cannot confirm causal relationships in this cross-sectional study, in which asthma could have led to methylation changes or vice versa.

In summary, we identified significant methylation-expression pairs in an eQTM analysis of nasal airway epithelium of subjects with and without asthma. Most methylation probes were associated with expression of distant cis-genes, and eQTM genes were enriched in immune regulation and epithelial integrity. Moreover, eQTM methylation probes and eQTM genes were over-represented among those associated with atopic asthma, further suggesting a key role of epigenetic regulation of gene expression in airway epithelium in disease pathogenesis.

## Methods

### Study population

Subject recruitment and study procedures for the Epigenetic Variation and Childhood Asthma in Puerto Ricans (EVA-PR) have been previously described^2^. In brief, EVA-PR is a case-control study of asthma in subjects aged 9-20 years. Participants with and without asthma were recruited from households in San Juan (PR) from February 2014 through May 2017, using multistage probability sampling; 638 households had ≥ 1 eligible subject, and 543 (85.1%) subjects (one per household) agreed to participate. There were no significant differences in age or sex between eligible children who did and did not participate. The study was approved by the institutional review boards of the University of Puerto Rico (San Juan, PR) and the University of Pittsburgh (Pittsburgh, PA). Written parental consent and assent were obtained from participants <18 years old, and consent was obtained from participants ≥ 18 years old.

The study protocol included questionnaires on respiratory health, measurement of serum allergen-specific IgEs, and collection of nasal epithelial samples for DNA and RNA extraction. Atopy was defined as ≥ 1 positive IgE (≥0.35 IU/mL) to five common allergens in Puerto Rico: house dust mite (Der p 1), cockroach (Bla g 2), cat dander (Fel d 1), dog dander (Can f 1), and mouse urinary protein (mus m 1). Asthma was defined as a physician’s diagnosis plus at least one episode of wheeze in the previous year. Control subjects had neither physician-diagnosed asthma nor wheeze in the previous year.

### Genome-wide study of DNA methylation and RNA sequencing

DNA and RNA were extracted from nasal specimens collected from the inferior turbinate. To account for potential effects of different cell types, we implemented a protocol in a subset of nasal samples (n=31) to select CD326-positive nasal epithelial cells before DNA and RNA extraction. Whole-genome methylation assays were done with HumanMethylation450 BeadChips (Illumina), as previously described^2^. Beta-values, ranging from 0 to 1, were calculated to measure percentage methylation at each CpG site. We then transformed beta values to M values because M values are closer to having a normal distribution (for linear regression analysis). As previously described, RNASeq was conducted with the Illumina NextSeq 500 platform (Illumina), paired-end reads at 75 cycles, and 80M reads/sample; reads were aligned to reference human genome (hg19) and transcripts per kilobase million (TPM) were used as proxy for gene expression level^2^. We excluded genes with low expression levels (mean TPM < 1) and genes whose transcription start site (TSS) was unavailable in hg19. TPM values were transformed to log_2_(TPM+1) for data analysis.

### eQTM analysis

We focused on identifying cis-eQTMs (i.e., CpGs regulating transcription of neighboring genes), due to limited power to perform a trans analysis (i.e., CpGs regulating distant genes)^23^. Thus, we only considered methylation probes within 1 Mb from the TSS of a gene. Using this criterion, we tested 8,552,964 methylation-gene expression pairs in analyses with and without adjustment for covariates. The unadjusted analysis was conducted to filter out potential false positive signals due to adjustment for batch effects^24,25^. Of the 24,171 methylation-expression pairs with a false-discovery rate-adjusted P <0.01 (FDR-P, see below) in the adjusted analysis, 7,304 pairs had an FDR-P ≥0.01 in the unadjusted analysis and were thus excluded from further consideration. Thus, we identified 16,867 methylation-expression pairs that were significant in both unadjusted and adjusted analyses.

For the adjusted analysis, we fitted a multivariate linear regression model; *y* = *β*_0_ + *β*_1_ *M*+ ***Tα*** + *ε*, where y is gene expression, *M* is methylation value at a probe, ***T*** represents other covariates, and *β*_0_, *β*_1_ and ***α*** are their regression coefficients. In this analysis, other covariates were asthma and atopy status, age, gender, the top five principal components from genotypic data, RNA sample sorting protocol (i.e., whole-cells or CD326-positive nasal epithelial cells), methylation and RNA-Seq batch, and latent factors that capture data heterogeneity from methylation and RNA-seq - estimated from R package sva ^26^. To conduct an efficient analysis, we used matrix eQTL package ^27^ to obtain P-values. FDR-P values were then calculated, based on all the methylation-expression pairs tested. For the unadjusted model, we only included methylation value as the following; *y* = *β*_0_ + *β*_1_ *M*+ *ε*.

### Epigenome-wide association study (EWAS) of atopic asthma

The EWAS conducted by Forno et al^2^ included 273 Puerto Rican subjects in EVA-PR (169 with atopic asthma and 104 control subjects without atopy or asthma). After quality controls, 227,836 methylation probes were evaluated in a multivariable logistic regression model, as follows: *logit(p)* = *β*_0_ + *β*_1_ *M*+ Σ *α*_*j*_*Z*_*j*_, where *p* is the probability of having atopic asthma, *M* is a methylation value at a probe, *Z*_*j*_ is an adjusted covariate, and *β*_0_, *β*_1,_ and *α*_*j*_ are regression coefficients. Other covariates included in the model were the first five PCs derived from genotypic data, age, gender, methylation batches, and latent factors of methylation –estimated from R package sva^26^. FDR-P values were then calculated based on testing 227,836 methylation probes. Significance was defined as FDR-P <0.01.

### Transcriptome-wide association studies (TWAS) of atopic asthma

A TWAS of atopic asthma was recently conducted by Forno et al^11^ in 258 Puerto Rican subjects in EVA-PR (157 with atopic asthma and 101 non-atopic non-asthmatic control subjects). In that study, differential gene expression was analyzed based on the raw count table of RNA sequencing data used using the R package DESeq2. Multivariable models of atopic asthma were adjusted for age, gender, RNA batches, RNA cell sorting, and the first five PCs derived from genotypic data. FDR-P were calculated for the 18,311 genes tested. Significance was defined as an FDR-P <0.01.

### Mediation analysis

To understand how methylation affects asthma through gene expression as a putative mediator, we conducted mediation analyses to identify indirectly associated methylation CpGs to atopic asthma through gene expression. We used the Baron and Kenny approach ^28^ instead of the Sobel method ^29^, due to differences in sample size between the eQTM analysis (including all subjects) and that for atopic asthma (including only subjects with atopic asthma and non-atopic controls).

To have a significant mediation of gene expression, all of the following needed to be significant: 1) the association between methylation and gene expression 2) the association between methylation and atopic asthma 3) the association between gene expression and atopic asthma. For #1, we only considered eQTM methylation probes and genes as candidates for the mediation tests. For #2, we recalculated FDR-P values of the result from our prior EWAS^2^ only for the eQTM probes, to reduce multiple testing. For #3, we conducted a TWAS fitting a logistic regression model: *logit(p)* = *β*_0_ + *β*_1_*X*+ Σ *α*_*j*_ *A*_*j*_, where *p* is the probability of having atopic asthma, *X* is a gene, *A*_*j*_ is an adjusted covariate, and *β*_0_, *β*_1,_ and *α*_*j*_ are regression coefficients. The adjusted covariates included in the model were the first five PCs derived from genotypic data, age, gender, whether RNA samples were from CD326-positive nasal epithelial cells, RNA batches, and a latent factor of gene expression, calculated from the R package sva^26^.

## Acknowledgements

This study was supported by grants HL079966, HL117191, MD011764 (to Juan C. Celedón), and U54MD007587 (to the University of Puerto Rico) from the U.S. National Institutes of Health (NIH). Dr. Kim is supported by a T32 training grant (HL129949) from the U.S. NIH. Dr. Forno’s contribution was supported by NIH grant HL125666. Dr. Yan’s contribution was supported by NIH grant K01 HL138098.

## Author Contributions

W.C. and J.C.C. conceived and designed the study. S.K. conducted the primary analysis and interpreted data. E.F., R.Z., Q.Y., N.B., E.A-P., and G.C. participated in data collection and data analysis. S.K., E.F., W.C. and J.C.C. prepared the first draft of the manuscript. All authors reviewed the draft for intellectual content, and approved submission of the final version of the manuscript.

## Data Availability

Datasets generated and analyzed during the current study are not publicly available because we did not obtain consent for such public release of epigenetic and transcriptomic data from participants. However, raw data to generate figures and tables are available from the corresponding author with the appropriate permission from the EVA-PR study team and the corresponding author upon reasonable request and institutional review board approval.

